# FUCCI tracking shows that Neurog3 levels vary with cell-cycle phase in endocrine-biased pancreatic progenitors

**DOI:** 10.1101/153700

**Authors:** Matthew E. Bechard, Eric D. Bankaitis, Alessandro Ustione, David W. Piston, Mark A. Magnuson, Christopher V.E. Wright

## Abstract

Neurog3^HI^ endocrine-committing cells are generated from a population of Sox9^+^ mitotic progenitors with only a low level of *Neurog3* transcriptional activity (*Neurog3*^TA.LO^). Low-level Neurog3 protein, in *Neurog3*^TA.LO^ cells, is required to maintain their mitotic endocrine-lineage-primed status. Herein, we describe a *Neurog3*-driven FUCCI cell-cycle reporter (*Neurog3*^P2A.FUCCI^) derived from a *Neurog3* BAC transgenic reporter that functions as a loxed cassette acceptor (LCA). In cycling Sox9^+^ Neurog3^TA.LO^ progenitors, the majority of cells in S-G_2_-M phases have undetectable levels of Neurog3 with increased expression of endocrine progenitor markers, while those in G_1_ have low Neurog3 levels with increased expression of endocrine differentiation markers. These findings support a model in which variations in Neurog3 protein levels are coordinated with cell-cycle phase progression in *Neurog3*^TA.LO^ progenitors with entrance into G_1_ triggering a concerted effort, beyond increasing Neurog3 levels, to maintain an endocrine-lineage-primed state by initiating expression of the downstream endocrine differentiation program prior to endocrine-commitment.

## Introduction

*Neurogenin3* (*Neurog3*) encodes a bHLH transcription factor essential for endocrine-lineage specification during mouse pancreas organogenesis (**1**). *Neurog3* is also critical to human pancreatic endocrine-cell development, with null mutations causing neonatal diabetes, and blocking β-cell differentiation from hESC (**2**). During mouse pancreatic development, high-level *Neurog3* expression (*Neurog3*^HI^) in Sox9^+^ pancreatic epithelial cells causes cell-cycle exit, endocrine commitment and epithelial delamination (**3–6**). We recently demonstrated, however, that low Neurog3 levels are necessary for maintaining a population of Sox9^+^ *Neurog3-* transcriptionally-active pancreatic epithelial cells in a mitotic endocrine-biased progenitor state (defined as *Neurog3*^TA.LO^), which pre-empts the transition to an endocrine-committed Neurog3^HI^ state (**6**,**7**). Our findings presented a significant parallel to how a low level of Neurog2 promotes a neural-progenitor state while high levels cause neural differentiation and cell-cycle exit (**8**–**11**). In those studies, higher Cdk activity in rapidly cycling progenitors, which have a relatively short G_1_, keeps Neurog2 in a (hyper)-phosphorylated, unstable state that activates neural-progenitor target genes (**10**,**12**). When the cell cycle of neural progenitors lengthens, however, and G_1_ lengthens, Cdk activity decreases, resulting in accumulation of a more stable (hypo)-phosphorylated Neurog2 that preferentially activates neural-differentiation targets (**10**,**12**). Recently, we demonstrated that keeping Neurog3 levels low leads to an increased mitotic index of *Neurog3*^TA.LO^ progenitors and expands their numbers within the pancreatic epithelium (**6**). Moreover, time-lapse observations show that the transition from the low level of Neurog3 observed in mitotic *Neurog3*^TA.LO^ progenitors to the high level necessary for endocrine-commitment occurs ~3-6 hours after division of the parental *Neurog3*^TA.LO^ cell, during G_1_ (**6**). These findings led to our proposal that the level and stability of Neurog3 in mitotic Sox9^+^ *Neurog3*^TA.LO^ progenitors is regulated by the cell cycle and that G_1_ extension promotes Neurog3 stabilization, accumulation, and endocrine commitment (**13**). Two recent reports support this model, demonstrating that Neurog3 is targeted and destabilized by Cdks and that G_1_ lengthening, by reducing Cdk activity, causes the accumulation of a more stable un(der)phosphorylated form of Neurog3 (**14**,**15**).

We have been independently investigating if Neurog3 protein stability and progenitor maintenance vs. endocrine differentiation decisions are connected to cell-cycle progression in *Neurog3*^TA.LO^ progenitors. To do so, we used recombinase-mediated cassette exchange (RMCE) to replace our previously described *Neurog3*^RG^ BAC transgenic reporter – which was designed as a Loxed Cassette Acceptor (LCA) – with a *Neurog3*-driven single-transgene insert of the FUCCI (*F*luorescence *U*biquitin *C*ell *C*ycle *I*ndicator) reporter (*Neurog3*^P2A.FUCCI^). Our analysis of *Neurog3*^P2A.FUCCI^ reporter activity showed that in cycling Sox9^+^ *Neurog3*^TA.LO^ progenitors, Neurog3 protein levels are highest during G_1_ and lowest during S-G_2_-M. Moreover, Sox9^+^ *Neurog3*^TA.LO^ progenitors in early G_1_ show increased expression of downstream Neurog3 targets usually associated with the forward passage into an endocrine commitment and progression program. We propose that these findings support a model in which the endocrine-differentiation program is already accessed, or preformed (albeit at a low or incomplete level), in mitotic *Neurog3*^TA.LO^ progenitors prior to moving into endocrine-commitment. This work provides a new tool for investigating, under *in vivo* conditions, Neurog3 and cell-cycle connections in lineage-primed progenitors, and new insight on the role of Neurog3 in regulating progenitor maintenance vs endocrine-commitment decisions.

## Results and Discussion

### Generating a Neurog3-driven P2A-fused single transgene FUCCI reporter

The FUCCI reporter relies on cell-cycle-phase-dependent destruction of fluorescent proteins fused to “degradation boxes” from hGeminin and hCdt, specifically the regions hGem^(1/110)^ and hCdt1^(30/120)^ (**16**), allowing cell-cycle phase determination (Figure 1A). To investigate connections between Neurog3 protein levels and cell-cycle progression in *Neurog3*^TA.LO^ progenitors we generated a single mKO2-hCdt1^(30/120)^-P2A-mVenus-hGem^(1/110)^ FUCCI (P2A.FUCCI) cassette, enabling both FUCCI components to be expressed under the control of *Neurog3* (Figure 1B). We selected the pairing of mKO2/mVenus because their fluorophores are spectrally separable from GFP and mCherry, allowing P2A.FUCCI visualization in cells carrying our previously described *Neurog3*- driven H2B^mCherry^-P2A-GFP^GPI^ (*Neurog3*^RG1^ reporter) (**6**). As seen with the original FUCCI reporter (**16**), *CMV*-driven expression of P2A.FUCCI in HeLa cells resulted in mKO2-hCdt1^(30/120)^ positivity during G_1_ and mVenus-hGem^(1/110)^ positivity during S-G_2_-M, with a brief overlap of the two fusion proteins during the G_1_/S phase transition (Figure 1B).

**Figure 1:**
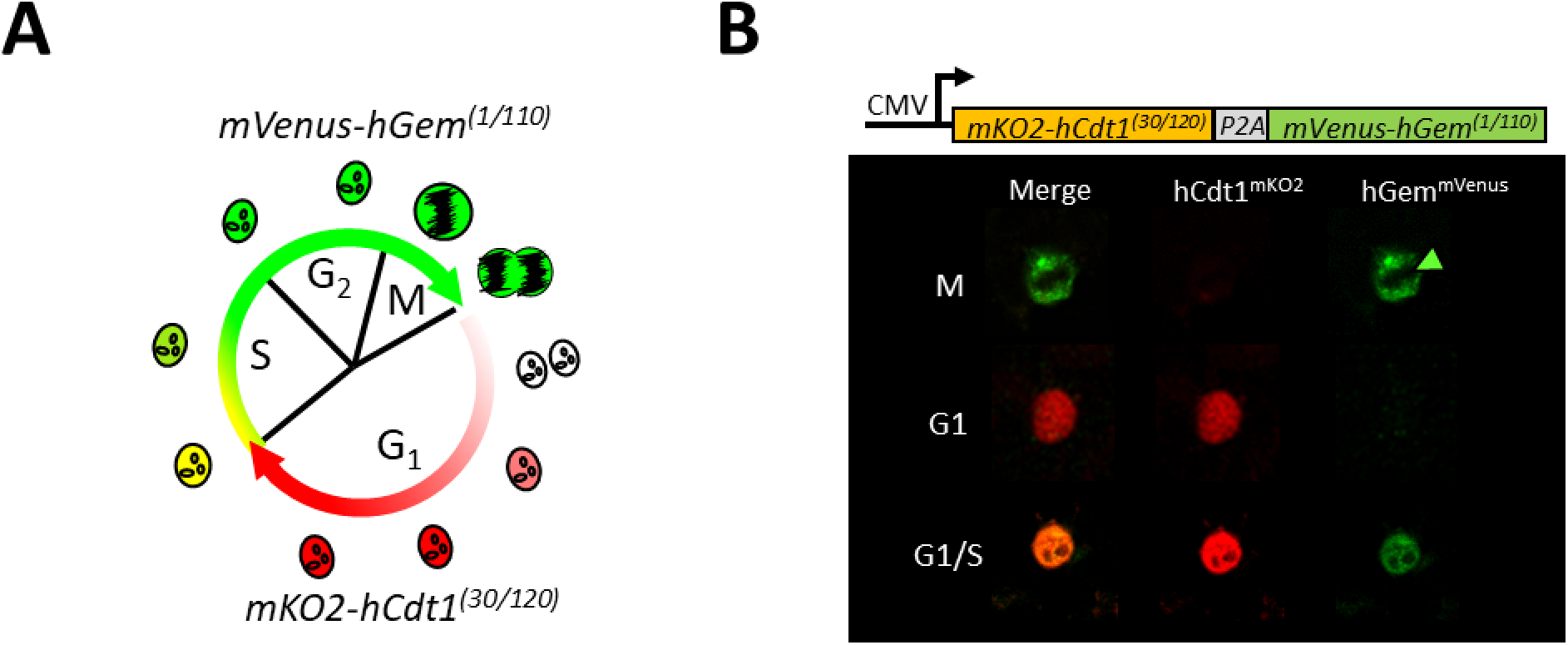
Peptide-2A single-transgene FUCCI transgene. (A) Diagram (adapted from Sakaue-Sawano et al., 2008) indicating phases of the cell cycle marked by the components of the FUCCI reporter: mVenus-hGem^(1/110)^ (S-G_2_-M) and mKO2-hCdt1^(30/120)^ (G_1_). (B) *Top*, Diagram of *CMV*^P2A.FUCCI^ expression plasmid. *Bottom*, Immunofluorescence images showing *CMV*^P2A.FUCCI^ reporter expression in HeLa cells at appropriate stages of the cell cycle. Green arrowhead indicates mitotic chromosomes.

To facilitate generating additional *Neurog3* BAC transgenic reporters from the *Neurog3*^RG^ reporter, we had flanked the *Neurog3*^RG^ cassette with tandem *lox71* and *lox2272* sites, making a Loxed Cassette Acceptor (LCA) allele (see ref. **6**; Figure 2A). This design was to allow *Neurog3*^RG^ to be replaced with any lox66/lox2272-flanked cassette via RMCE in mESCs. To avoid potential issues with performing RMCE in cells carrying multiple LCA alleles, mESCs identified as having stably integrated the *Neurog3*^RG^ BAC LCA transgene were screened for single-copy insertion by a qPCR-based assay (see methods and materials; Figure 2B and C) that accurately estimates transgene copy number (**17**). Using this assay, two mESC transgenic clones, referred to as *Neurog3*^RG1^ and *Neurog3*^RG2^, were identified as having copy numbers of 1.25 ±0.16 and 1.46 ±0.26 (Figure 2C). Derivation of *Neurog3*^RG^ mESC lines and the subsequent *Neurog3*^RG1^ mouse line are described in (**6**). As with the *Neurog3*^RG1^ mouse (**6**), examination, in *Neurog3*^RG2^ mice, of gross tissue and islet architecture, ad libitum fed glucose levels, and proportions of Sox9^+^ Neurog3 protein-low (Neurog3^pLO^) versus Sox9^-^ Neurog3 protein-high (Neurog3^pHI^) cells during pancreas development, revealed no abnormal phenotype (Figure 2-figure supplement 1A-D; data not shown). We next validated the LCA function of the *Neurog3*^RG^ BAC transgene by using the *Neurog3*^RG2^ mESC line to derive a *Neurog3*^P2A.FUCCI^ mESC line. A lox66/lox2272-flanked *Neurog3*^P2A.FUCCI^-PGK-hygro^R^ cassette was generated with cassette placement mimicking that of *Neurog3*^RG^ (Figure 2-figure supplement 2A). Following RMCE in *Neurog3*^RG2^ mESCs, PCR was performed to verify replacement of the lox71/lox2272-flanked *Neurog3*^RG^-PGK-Puro^ΔTK^ cassette with the lox66/lox2272 *Neurog3*^P2A.FUCCI^-PGK-hygro^R^ cassette (Figure 2-figure supplement 2B). This derivative *Neurog3*^P2A.FUCCI^ mESC line was then used to generate *Neurog3*^P2A.FUCCI^ transgenic mice. Given that the genomic integration site is likely different in *Neurog3*^RG2^ vs. *Neurog3*^RG1^ mESC lines, we used *Neurog3*^RG2^ mESCs to derive *Neurog3*^P2A.FUCCI^ mice to allow future breeding of *Neurog3*^P2A.FUCCI^ to *Neurog3*^RG1^ mice to enable four-color reporting of cell-cycle phase and *Neurog3* expression. Our proposal is that such visualization could facilitate experiments aimed at understanding if, like other progenitor populations, G_1_ length or overall cell-cycle length in mitotic *Neurog3*^TA.LO^ progenitors plays a role in regulating progenitor maintenance vs. endocrine-commitment decisions, or even in determining whether one endocrine cell-type is produced over another at specific stages or locations within the developing pancreas.

**Figure 2:**
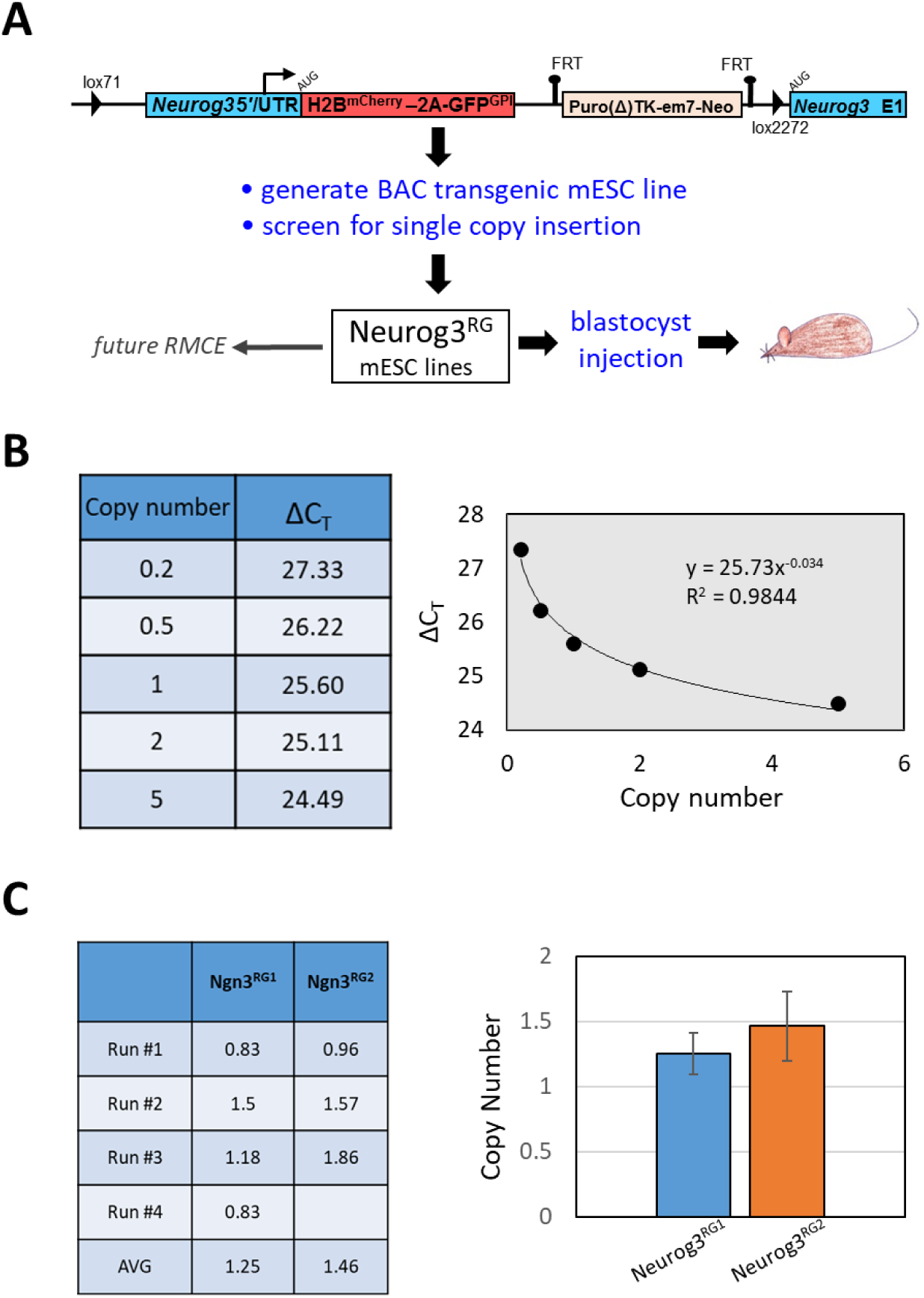
Generation of an LCA-capable BAC transgenic *Neurog3*^RG^ mESC line. (A) Schematic detailing the generation of transgenic mES cell lines carrying a single copy of a *Neurog3*^RG^ BAC transgene designed to serve as an LCA in future RMCE reactions. The *Neurog3*^RG^ BAC transgenic mESCs were previously used to generate *Neurog3*^RG^ reporter mice (**6**). Neurog3 5′/UTR represents the region 5′ of the start codon containing cis regulatory elements and the Neurog3 5′ untranslated region (UTR). (B) Table and graph of a standard curve, generated via a qPCR-based assay (see methods and materials), that relates transgene copy number to a specific ΔC_T_ value. (C) Table and graph depicting the estimated *Neurog3*^RG^ BAC transgene copy number present in *Neurog3*^RG1^ and *Neurog3*^RG2^ mESC lines.

### Neurog3 levels and progenitor maintenance vs. endocrine-commitment decisions are coupled to the cell cycle

Recent studies showed that during S-G_2_-M, Neurog3 is kept in a hyperphosphorylated unstable state via Cdk phosphorylation, and that decreased Cdk activity associated with entrance into G_1_ result in stabilization and accumulation (**14**,**15**). These findings support our model that, in actively cycling *Neurog3*^TA.LO^ progenitors, Neurog3 protein levels vary according to the cell-cycle phase (**13**). To address this issue, we used *Neurog3*^P2A.FUCCI^ reporter expression in *Neurog3*^TA.LO^ progenitors to track cell-cycle progression in relation to Neurog3 protein levels. Previously, *Neurog3*^TA.LO^ progenitors were defined as a population of Sox9^+^ *Neurog3*-transciptionally active (low-level *Neurog3*^RG1^ reporter expression) progenitors comprising cells with either low (*Neurog3*^TA.pLO^) or immunologically undetectable Neurog3 (*Neurog3*^TA.pUD^) (**6**). Consistent with this definition, *Neurog3*^TA.LO^ progenitors were herein defined as Sox9-positive and positive for either component of the *Neurog3*^P2A.FUCCI^ reporter, with low or undetectable Neurog3 protein, whereas endocrine-committed *Neurog3*^TA.HI^ cells should be Sox9-negative, Neurog3^pHI^ and positive for mKO2-hCdt1^(30/120)^ (Figure 3A). Unexpectedly, we detected significant residual cytoplasmic mVenus-hGem^(1/110)^ fluorescence in post-mitotic, actively delaminating, endocrine-committed *Neurog3*^TA.HI^ cells that showed the expected high mKO2-hCdt1^(30/120)^ signal (Figure 3A). This observation was different from previous reports on the FUCCI reporter, where mVenus-hGem^(1/110)^ was mostly degraded after M-phase, becoming absent by the time of mKO2-hCdt^(30/120)^ detection in early G_1_ (**16**,**18**). This cytoplasmic mVenus-hGem^(1/110)^ signal, however, was completely absent in islets (data not shown), which could suggest that high *Neurog3*^P2A.FUCCI^ reporter expression in delaminating *Neurog3*^TA.HI^ cells overwhelms the ubiquitin-mediated protein degradation pathway, extending the time necessary to fully degrade mVenus-hGem^(1/110)^ after entering G_1_. We were therefore careful to score *Neurog3*^P2A.FUCCI^ cells as only in S-G_2_-M if definitively nuclear mVenus signal was observed, with no indication of mKO2 (Figure 3A). By these criteria, the majority of Sox9^+^ *Neurog3*^TA.LO^ cells were in S-G_2_-M and thus mitotic, while nearly all *Neurog3*^TA.HI^ cells were in G_1_ (Figure 3B). To determine if Neurog3 protein levels vary through the cell cycle, we examined the cell-cycle status of Sox9-positive *Neurog3*^TA.pLO^ versus *Neurog3*^TA.pUD^ cells. Quantification revealed that 78% (± 8.1%) of *Neurog3*^TA.pUD^ cells were in S-G_2_-M and 22% (± 8.1%) in G_1_ (Figure 3B). Although stabilization and accumulation of Neurog3 occurs during G_1_, previous work showed that Neurog3 protein is present during S phase, with rapid degradation occurring during G_2_-M (**14**). Corroborating that result, we show that while the majority (55% ± 3.6%) of Sox9^+^ *Neurog3*^TA.pLO^ cells were in G_1_, 45% (± 3.6%) were in S-G_2_-M (Figure 3B). These findings show that the Neurog3 protein level in cycling Sox9^+^ *Neurog3*^TA.LO^ progenitors is lowest during S-G_2_-M and highest during G_1_.

**Figure 3:**
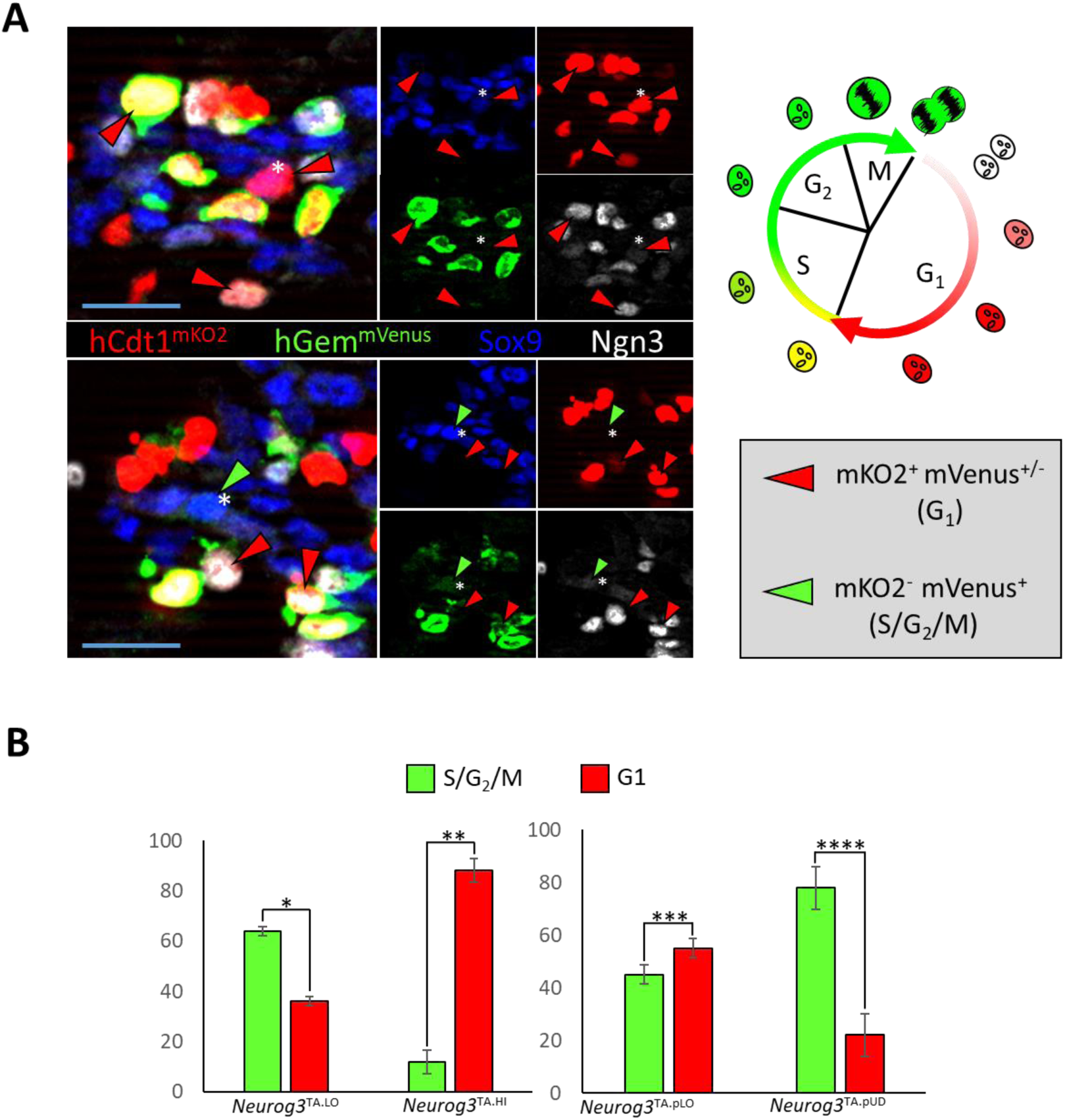
Neurog3 protein levels vary according to cell-cycle phase. (A) E14.5 pancreatic epithelium showing Sox9, Neurog3, hCdt1^mKO2^ and hGem^mVenus^. Red arrowheads indicate *Neurog3*^P2A.FUCCI+^ cells that are mKO2^+^ and thus in G_1_, green arrowheads indicate *Neurog3*^P2A.FUCCI+^ cells that are mVenus^+^ and thus in S-G_2_-M phase. Asterisk indicates Sox9^+^ *Neurog3*^TA.LO^ cells, arrows with no asterisk indicate Sox9^-^ *Neurog3*^TA.HI^ cells. (B) Left, percentage of Sox9^+^ *Neurog3*^TA.LO^ and Sox9^-^ *Neurog3*^TA.HI^ cells in S-G_2_-M versus G_1_ phase. Right, percentage of Sox9^+^ *Neurog3*^TA.pLO^ and Sox9^+^ *Neurog3*^TA.pUD^ in S-G_2_-M versus G_1_ phase. (*n* = 1600, *N* = 3). (*) *P* = 0.0072; (**) *P* = 4 x 10^-6^; (***) *P* = 0.0607; (****) *P* = 0.0039. Data are mean ± SEM. Bars, 20 μm.

Previous work shows that a low Neurog3 protein level maintains a mitotic, endocrine lineage-primed progenitor state (**1**,**4**–**7**). Given the cell-cycle-dependent variation of Neurog3 protein level, we hypothesized that the low-level accumulation of Neurog3 in *Neurog3*^TA.LO^ progenitors in G_1_ could trigger gene expression changes that were consistent with endocrine lineage-priming, and involving genes other than solely *Neurog3*. Therefore, intraepithelial Neurog3^P2A.FUCCI+^ cells (*Neurog3*^TA.LO^ progenitors) were isolated from E14.5 *Neurog3*^P2A.FUCCI^ pancreatic explants by flow sorting of lumen-contacting (Muc1^+^) cells, then sorting cells in S-G_2_-M (mKO2^-^ mVenus^+^) or G_1_ (mKO2^+^ mVenus^-^) (Figure 4A). As described above, actively delaminating *Neurog3*^TA.HI^ cells display relatively bright, yet to be degraded, cytoplasmic mVenus and high nuclear mKO2 (Figure 3A). To exclude this population, the flow-cytometry gating was set so that mVenus/mKO2 co-positive cells were not collected (Figure 4A). Analysis via qRT-PCR showed that while in S-G_2_-M, *Neurog3*^TA.LO^ progenitors are enriched for *Sox9* and *Hes1* (mitotic endocrine-progenitor markers) with low expression of *Neurog3* and several markers indicating forward progression towards endocrine commitment and further differentiation *(NeuroD1*, *Insm1*, *Glucagon*, *Insulin*) (Figure 4B). *Neurog3*^TA.LO^ progenitors in G_1_, however, showed significantly decreased *Hes1* and increased *Neurog3*, *NeuroD1*, *Insm1*, *Glucagon* and *Insulin* (Figure 4B). Despite the increases in endocrine-commitment markers, entrance into G_1_ did not significantly alter *Sox9* expression (Figure 4B), demonstrating that these cells are intraepithelial *Neurog3*^TA.LO^ progenitors. The data are also consistent with the idea that cells in this mitotic progenitor state, when in G_1_, initiate expression of several genes representing the downstream endocrine differentiation program, at a stage prior to commitment. Given the role of Neurog3 in trans-activating *NeuroD1* and *Insm1* (**19**–**21**), we speculate that the low-level accumulation of Neurog3 specifically in G_1_ could be sufficient to induce low-level *NeuroD1/Insm1* expression in lineage-primed progenitors. It is also possible that signals initiating the lineage-primed state activate low-level expression of other transcription-factor genes in a Neurog3-independent manner. It is plausible that the concerted expression of several trans-acting factors establishes a relatively weak or incomplete form of the GRN that is normally considered to work only in post-mitotic committed cells. It would be important to discover if entrance into G_1_ were also linked to alterations in chromatin architecture and DNA accessibility that allow low-level expression of GRN member genes contributing to lineage priming in *Neurog3*^TA.LO^ progenitors. Understanding how cell-cycle progression regulates such gene expression programs could lead to understanding if the final hormone-secreting cell fate might become preconditioned in the mitotic lineage-biased stage, and possibly how to manipulate cells at this early phase of their lifespan to improve the generation of functional endocrine cells.

**Figure 4:**
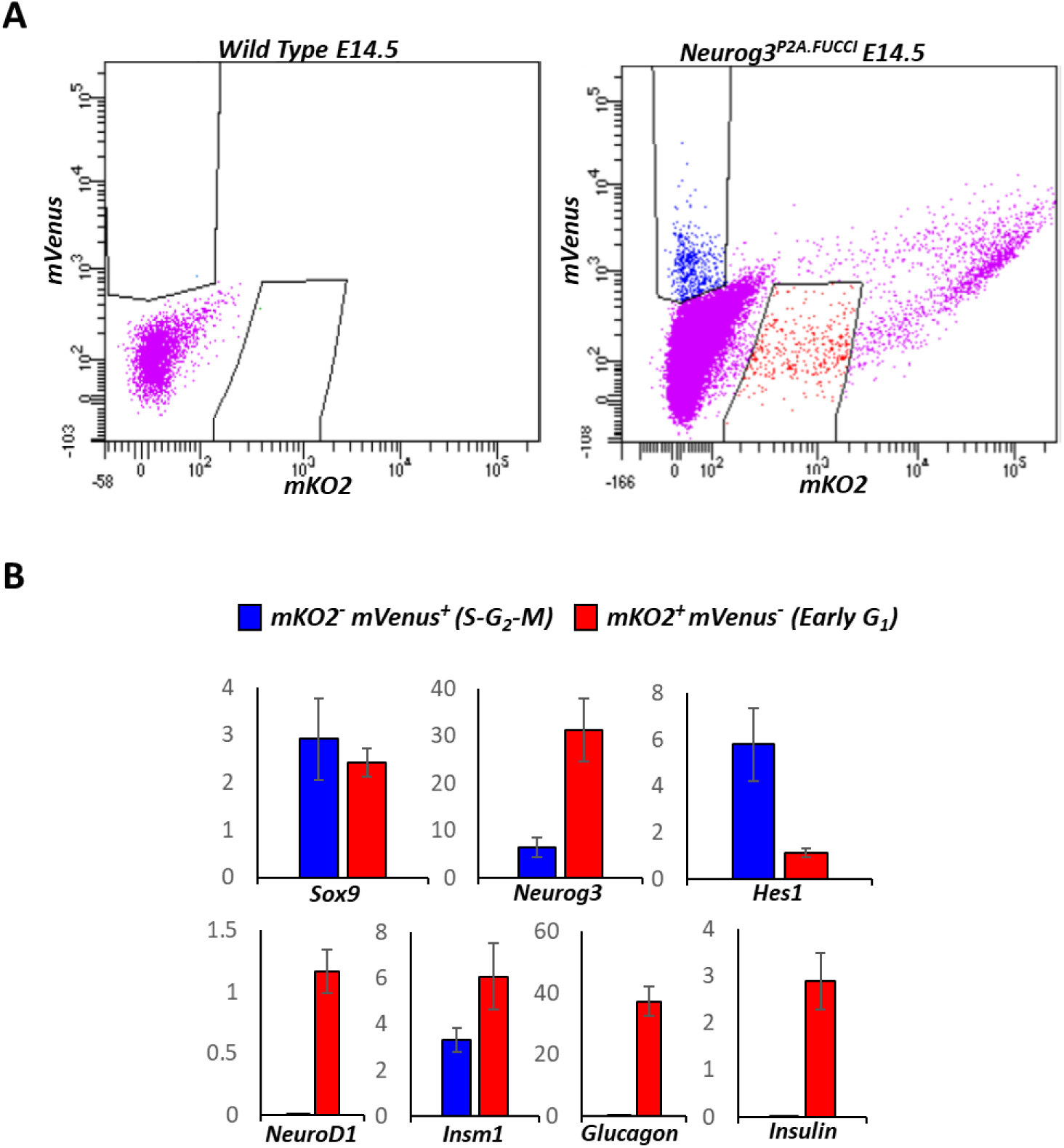
Neurog3 promotes low-level activation of downstream targets during G_1_ in the mitotic *Neurog3*^TA.LO^ progenitor state. (A) flow cytometry plot detailing capture of lumen-apposed (Muc1^+^) Neurog3^P2A.FUCCI+^ cells in S-G_2_-M (mKO2^-^ mVenus^+^) (Blue population) or G_1_ (mKO2^+^ mVenus^-^) (red population) from E14.5 *Neurog3*^P2A.FUCCI^ pancreata. Flow-sorted cells were collected into TRIzol for RNA isolation and cDNA synthesis. (B) Relative expression level (*y*-axis), normalized to *Gapdh*, of *Sox9*, *Neurog3*, *Hes1, NeuroD1, Insm1, glucagon*, and *Insulin* for E14.5 flow captured Muc1^+^ *Neurog3*^P2A.FUCCI+^ mKO2^-^ mVenus^+^ (blue bars) and Muc1^+^ *Neurog3*^P2A.FUCCI^ mKO2^+^ mVenus^-^ (red bars) cells. Each data point represents an average of at least three technical replicates. Error bars are SEM. See supplemental table 1 for a list of primers used.

## Materials and Methods

### Mice and transgene copy number analysis

Animal protocols were approved by the Vanderbilt University Institutional Animal Care and Use Committee. All animals were PCR genotyped. Sequences for genotyping primers are listed in Supplemental Table 1. Generation of the *Neurog3*^RG^ BAC transgene and subsequent derivation of the *Neurog3*^RG2^ mESC line and *Neurog3*^RG2^ reporter mice was described previously (**6**). Although not previously reported in (**6**), mouse ES cells that stably integrated the *Neurog3*^RG^ BAC LCA were analyzed by a qPCR-based assay that accurately estimates transgene copy number (**17**). Briefly, we generated primers specific for the puromycin-resistance gene (Puro^R^) in the Puro^R^-ΔTK-em7-Neo^R^ (Puro^ΔTK^) selection cassette in the *Neurog3*^RG^ transgene (Figure 2A). Quantitative PCR was run on 2.5, 10, 20, 40 and 200 ng of genomic DNA from a TL1 mESC knock-in line, carrying one copy of the Puro^ΔTK^ cassette inserted via homologous recombination, to yield a ΔC_T_ curve reflecting copy number (Figure 2B). Triplicate runs of exactly 20 ng of DNA from 23 candidate *Neurog3*^RG^ mESC lines used the standard curve to define copy number.

### Generation of P2A.FUCCI transgene and the *Neurog3*^P2A.FUCCI^ reporter mouse line

To generate the mKO2-hCdt1^(30/120)^-P2A-mVenus-hGem^(1/110)^ (P2A.FUCCI) cassette, mKO2-hCdt1^(30/120)^ and mVenus-hGem^(1/110)^ were PCR-amplified from plasmids provided by Dr. Atsushi Miyawaki (RIKEN Brain Science Institute) (Sakaue-Sawano et al. 2008). Amplification of mKO2-hCdt1^(30/120)^ involved attaching a 40 bp *Neurog3* homology region 5′ of the mKO2 start codon along with the first 25 base pairs of a P2A sequence 3′ of mKO2. Amplification of mVenus-hGem^(1/110)^ involved attaching a 5′ BamHI site and a 3′ ApaI site. A third PCR was used to generate a P2A cassette with 25 base pairs of the 3′ end of mKO2-hCdt1^(30/120)^ attached to its 5′ end and a BamH1 site at its 3′ end. The resulting mKO2-hCdt1^(30/120)^ and P2A amplicons were then fused together by overlap extension PCR (**22**), using a forward primer specific for the mKO2-hCdt1^(30/120)^ amplicon and a reverse primer specific for the P2A amplicon. The resulting mKO2-hCdt1^(30/120)^-P2A amplicon was attached to the mVenus-hGem^(1/110)^ amplicon via the BamHI site and inserted into a pBS KS(-) vector. The resulting P2A.FUCCI cassette was removed from pBS KS (-) and inserted into a pCMV5 vector with a PGK-neomycin selection cassette for expression in HeLa cells (described below). The P2A.FUCCI cassette was also inserted in place of the RG cassette in the PL451-RG-FRT-Puro^R^-ΔTK-em7-Neo^R^-FRT-lox2272 vector described previously (**6**). Using BAC recombineering the resulting P2A.FUCCI-FRT-Puro^R^-ΔTK-em7-Neo^R^-FRT-lox2272 cassette was inserted immediately upstream of the Neurog3 start codon in the Neurog3-containing RPCI-23-121F10 BAC (**6**). Using BAC recombineering, the P2A.FUCCI-FRT-Puro^R^-ΔTK-em7-Neo^R^-FRT-lox2272 cassette was retrieved into a vector containing a lox66 site in a manner that ensured that placement of the lox66 site precisely mimicked that of its lox71 counterpart in the lox71/lox2272 flanked *Neurog3*^RG^ BAC LCA. Subsequently, the FRT-flanked Puro^R^-ΔTK-em7-Neo^R^ cassette was replaced with an FRT-flanked PGK-Hygro^R^ selection cassette. This final lox66/lox2272 flanked *Neurog3*^P2A.FUCCI^ exchange plasmid was linearized and used to replace, via RMCE, the *Neurog3*^RG^ BAC LCA in the *Neurog3*^RG2^ mESC line. Successful replacement with *Neurog3*^P2A.FUCCI^ was verified by PCR (Figure Supplement 2B). A single, verified, *Neurog3*^P2A.FUCCI^ mES cell line was expanded, karyotyped and injected into blastocyst-stage embryos to derive the *Neurog3*^P2A.FUCCI^ reporter mouse strain. The LCA capability of the *Neurog3*^RG1^ mESC line was also tested and shown to allow efficient RMCE of lox66/lox2272 flanked cassettes (data not shown).

### Cell culture

HeLa cells were cultured on tissue culture grade plastic at 37° C in Dulbecco’s Modified Eagle Medium (DMEM) supplemented with 10% fetal bovine serum (FBS), and 100 U/mL penicillin-streptomycin. Cells were passaged by adding 0.05% trypsin-EDTA to a plate of semi-confluent (<90%) cells. To test expression of the P2A.FUCCI reporter, HeLa cells were transiently transfected with the *CMV*^P2A.FUCCI^ expression plasmid using Lipofectamine 2000 (Thermo Fisher) according to manufacturer’s instructions. The same conclusions were obtained with stable clonal lines expressing *CMV*^P2A.FUCCI^ selected for neomycin resistance over 14 days (not shown).

### Immunodetection

E14.5 dorsal pancreata were fixed in 4% paraformaldehyde (4 hrs, 4°C) then equilibrated in 30% sucrose overnight at 4°C). A Leica CM3050S was used sucrose-equilibrated, OCT-embedded tissue (Tissue-Tek) into 10 μm tissue sections, sequentially placed on three separate sets of slides, each covering ^~^33% of the dorsal pancreas. Primary and Secondary antibodies are listed in Supplemental Table 2. All images are epifluorescence from a Zeiss ApoTome microscope with Zeiss Axiovision software.

### Flow sorting and qRT-PCR analysis

Multiple E14.5 *Neurog3*^P2A.FUCCI+^ dorsal pancreata were pooled and dispersed into a single-cell suspension using Accumax (Sigma) (protocol available on request). Dispersed samples were washed and incubated on ice, first with Muc1 antibody for 1 hr, then anti-hamster Cy5 secondary antibody for an additional hour. DAPI was added to ensure sorting of viable cells. Flow sorting used a BD FACSAria III. cDNA was generated using iScript cDNA synthesis kit (BioRad) from RNA isolated from flow-sorted cells after TRIzol extraction. PCR was performed in a Bio-Rad CFX96 with SsoFast EvaGreen Supermix (Bio-Rad) using at least three technical replicates. Relative expression level (normalized to *Gapdh*) was calculated by first assessing the ΔC_T_ between the gene of interest and *Gapdh* before converting the ΔC_T_ to relative expression level (2^ΔC^_T_). The results in Figure 4 were independently repeated (biological replicate) with similar results. Primer sequences, except for *Insm1* primers (Applied Biosystems), are listed in Supplemental Table 1.

### Quantification and statistics

Cell counting and fluorescence intensity quantifications were done using NIH ImageJ software. For quantifications “n” indicates total cells counted, with “N” number of individual dorsal pancreata analyzed. As previously stated approximately 33% of an entire dorsal pancreas was analyzed for each dorsal pancreas. Previous reports indicate that only 2% of the total pancreas volume needs to be systematically sampled and analyzed to obtain a relative error of ≤ 10% (**23**). Error bars generated using standard error of the mean (SEM), with Student’s t-test (one-tailed) used to calculate *p* values. *p* values were deemed significant when ≤ 0.05.

## Acknowledgements

We thank Atsushi Miyawaki (RIKEN Brain Science Institute) for the mKO2-hCdt1^(30/120)^/pCSII-EFMCS and mVenus-hGem^(1/110)^/pCSII-EF-MCS plasmids. This work utilized the Cell Imaging Shared Resource and Transgenic/ES Cell Shared Resource core facilities of the Vanderbilt Diabetes Research and Training Center funded by NIDDK grant 020593. Flow cytometry was performed in the VUMC Flow Cytometry Shared Resource supported by the Vanderbilt-Ingram Cancer Center (P30 CA68485) and the Vanderbilt Digestive Disease Research Center (DK0558404). Generation of *Neurog3*^RG2^ and *Neurog3*^P2A.FUCCI^ mice was supported in part by the Beta Cell Biology Consortium Mouse ES Cell Core funded by the NIDDK (U01DK072473). We thank Anna Means, Guoqiang Gu, and members of the Wright/Gu labs for discussions. This study was supported by the NIH/NIDDK (U01DK089570) and an American Heart Association fellowship to MB (13POST14240011).

---

**Figure Supplement 1:**
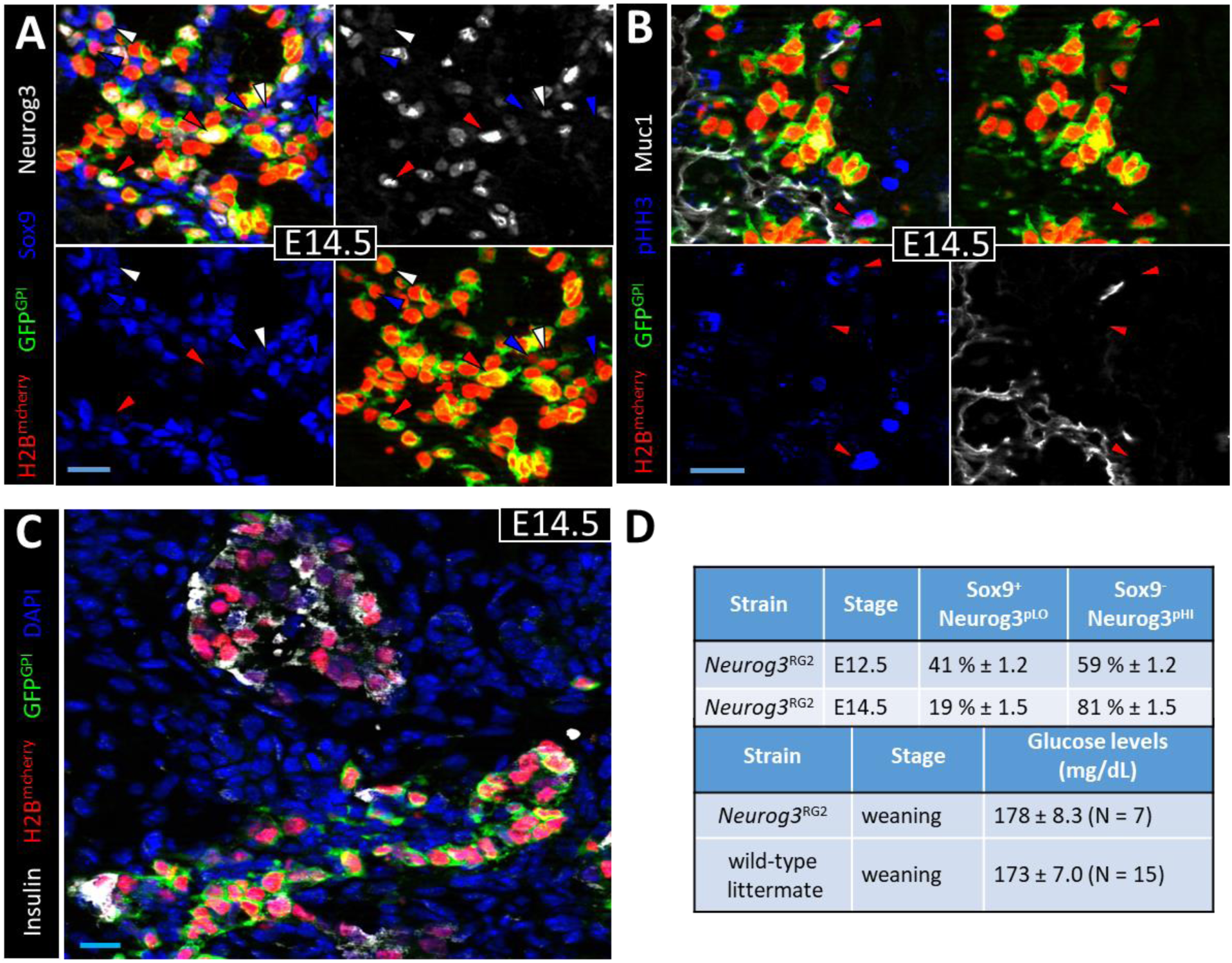
*Neurog3*^RG2^ BAC transgenic reporter is a passive reporter. (A) E14.5 pancreatic epithelium showing H2B^mCherry^, GFP^GPI^, Sox9 and Neurog3. Blue, white and red arrowheads indicate Sox9^+^ *Neurog3*^TA.pUD^ *cells*, Sox9^+^ *Neurog3*^TA.pLO^ cells and Sox9^-^ *Neurog3*^TA.HI^ cells, respectively. (B) E14.5 pancreatic epithelium showing H2B^mCherry^, GFP^GPI^, Muc1, and phospho-Histone H3 (pHH3). Red arrowheads indicate pHH3^+^ Muc1^+^ *Neurog3*^TA.LO^ cells. (C) Image of Islets of Langerhans and the pancreatic epithelium at E14.5 showing H2B^mCherry^, GFP^GPI^, Insulin and DAPI. (D) Top table details percentage of Sox9^+^ Neurog3^pLO^ vs. Sox9^-^ Neurog3^pHI^ cells in *Neurog3*^RG2+^ pancreatic epithelium at e12.5 and e14.5, which are unchanged relative to typical analyses of pancreata from wild-type mice (**5**). Bottom table details blood glucose levels, measured in whole blood using a Nova Max Plus glucose meter and test strips, of ad libitum fed wild-type and *Neurog3*^RG2+^ mice at weaning (~P21). Bars, 20 μm.

**Figure Supplement 2:**
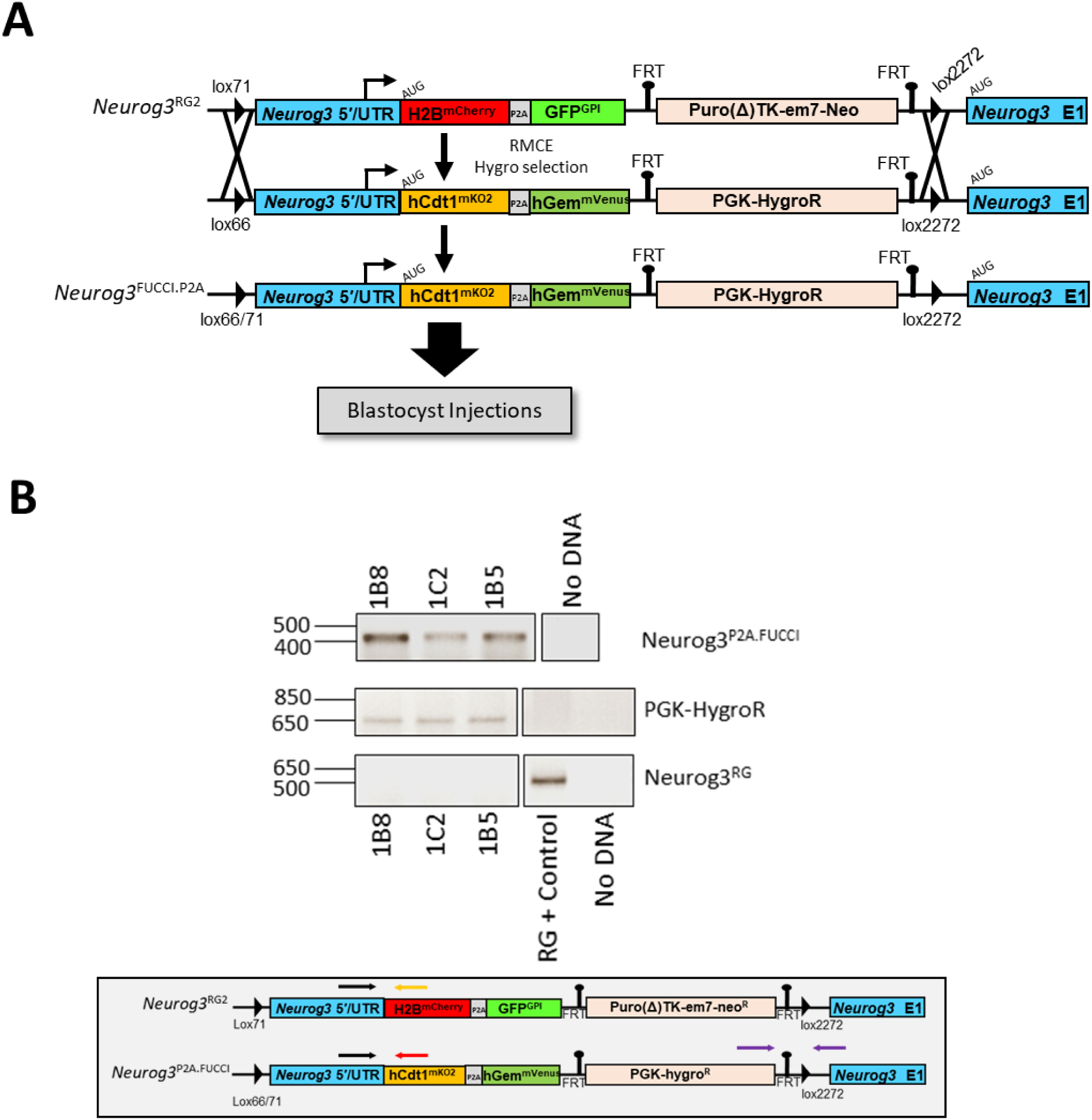
RMCE-mediated derivation of *Neurog3*^P2A.FUCCI^ mESC line. (A) Scheme for using RMCE to replace the LCA-capable *Neurog3*^RG^ BAC transgenic reporter with the Lox66/lox2272-flanked *Neurog3*^P2A.FUCCI^ transgenic reporter. (B) Top, PCR of genomic DNA from three mES cell lines (1B8, 1C2, 1B5) to check for successful RMCE of the lox71/lox2272-flanked *Neurog3*^RG^ cassette in *Neurog3*^RG2^ mESCs for the loxx66/lox2272-flanked *Neurog3*^P2A.FUCCI^ cassette. Bottom, schematic showing the approximate binding sites for the *Neurog3*^RG^, *Neruog3*^P2A.FUCCI^ and PGK-hygro^R^ primer pairs.

**Supplemental Table 1.**
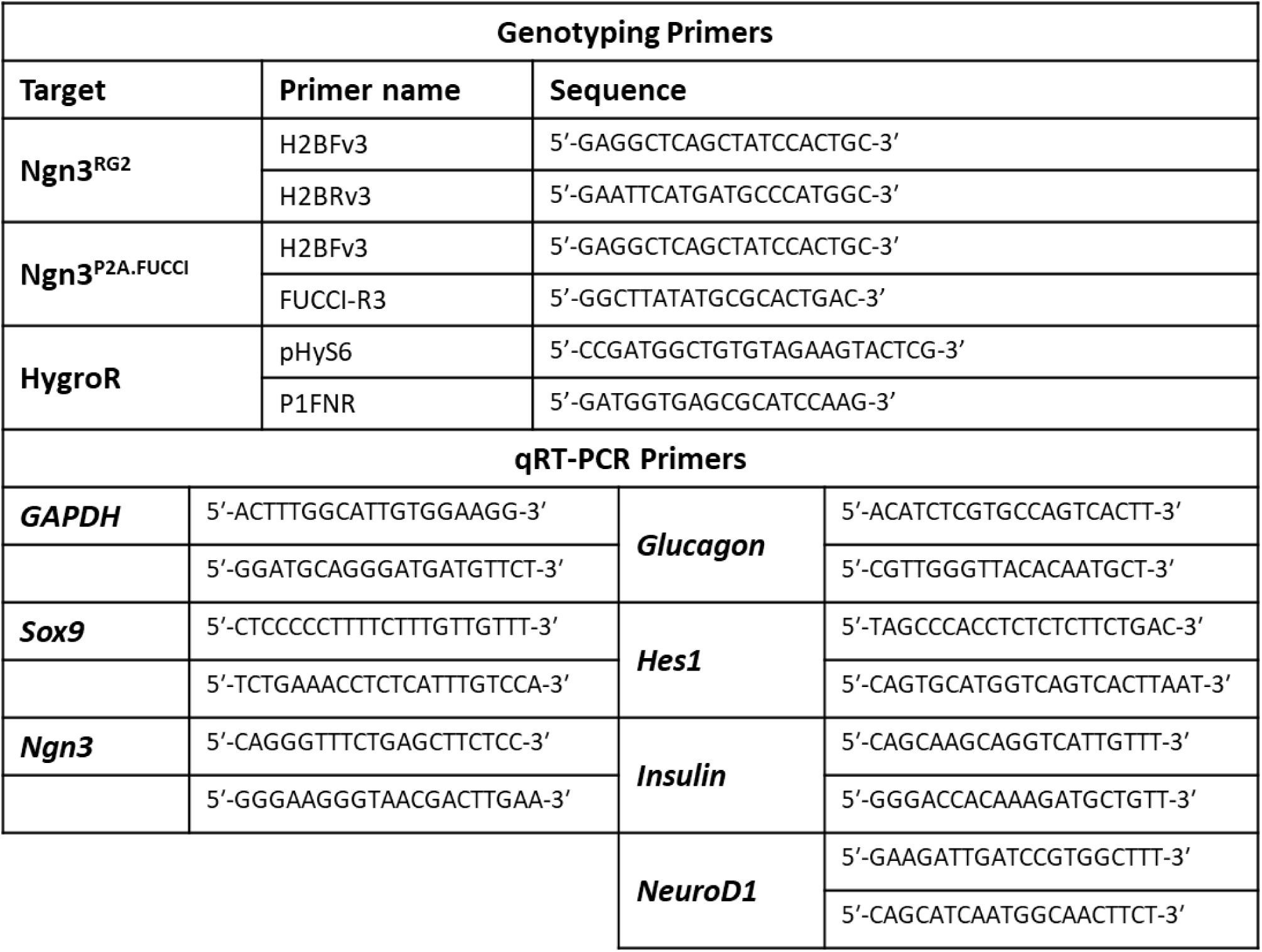
Primers used for genotyping and qRT-PCR analyses.

**Supplemental Table 2.** Antibodies and detection methods.

| Primary Antibodies |  |  |  |  |
| --- | --- | --- | --- | --- |
| Antigen | Species | Dilution | Label Method | Source |
| Muc1 | Hamster | 1:250 | IF | NeoMarkers |
| Sox9 | Rabbit | 1:2500 | IF | Millipore |
| Neurog3 | Goat | 1:40,000 | IF | G. Gu (Vanderbilt) |
| pHH3 | Rabbit | 1:500 | IF | Millipore |
| EGFP/Venus | Rabbit<br>Chicken | 1:500<br>1:500 | IF<br>IF | Clontech<br>Aves |
| Insulin | Guinea Pig | 1:500 | IF | Dako |
| DAPI | ---- | ---- | Mount Media | Life Technologies |
| Secondary Antibodies |  |  |  |  |
| Antigen | Conjugation | Dilution | Source |  |
| Rabbit/Chicken | Cy2 | 1:250 | Jackson ImmunoResearch |  |
| Guinea pig/ Goat<br>Hamster | Cy5 | 1:250 | Jackson ImmunoResearch |  |
| Rabbit | 405 | 1:250 |  |  |
| Goat | Biotinylated | 1:250 | Vector Laboratories |  |

## References

1. Gradwohl G, Dierich A, LeMeur M, Guillemot F. 2000. Neurogenin3 Is Required for the Development of the Four Endocrine Cell Lineages of the Pancreas. Proc Natl Acad Sci U S A 97:1607–1611.

2. McGrath PS, Watson CL, Ingram C, Helmrath MA, Wells JM. 2015. The Basic Helix-Loop-Helix Transcription Factor NEUROG3 Is Required for Development of the Human Endocrine Pancreas. Diabetes 64:2497–505.

3. Miyatsuka T, Kosaka Y, Kim H, German MS. 2011. Neurogenin3 inhibits proliferation in endocrine progenitors by inducing Cdkn1a. Proc Natl Acad Sci 108:185–190.

4. Wang S, Yan J, Anderson DA, Xu Y, Kanal MC, Cao Z, Wright CVE, Gu G. 2010. Neurog3 gene dosage regulates allocation of endocrine and exocrine cell fates in the developing mouse pancreas. Dev Biol 339:26–37.

5. Johansson KA, Dursun U, Jordan N, Gu G, Beermann F, Gradwohl G, Grapin-Botton A. 2007. Temporal Control of Neurogenin3 Activity in Pancreas Progenitors Reveals Competence Windows for the Generation of Different Endocrine Cell Types. Dev Cell 12:457–465.

6. Bechard ME, Bankaitis ED, Hipkens SB, Ustione A, Piston DW, Yang Y-P, Magnuson MA, Wright CVE. 2016. Precommitment low-level Neurog3 expression defines a long-lived mitotic endocrine-biased progenitor pool that drives production of endocrine-committed cells. Genes Dev 30:1852–1865.

7. Bankaitis ED, Bechard ME, Wright CVE. 2015. Feedback control of growth, differentiation, and morphogenesis of pancreatic endocrine progenitors in an epithelial plexus niche. Genes Dev 29:2203–2216.

8. Roybon L, Hjalt T, Stott S, Guillemot F, Li J-Y, Brundin P. 2009. Neurogenin2 directs granule neuroblast production and amplification while NeuroD1 specifies neuronal fate during hippocampal neurogenesis. PLoS One 4:e4779.

9. Shimojo H, Ohtsuka T, Kageyama R. 2011. Dynamic Expression of Notch Signaling Genes in Neural Stem/Progenitor Cells. Front Neurosci 5:1–7.

10. Ali F, Hindley C, McDowell G, Deibler R, Jones A, Kirschner M, Guillemot F, Philpott A. 2011. Cell cycle-regulated multi-site phosphorylation of Neurogenin 2 coordinates cell cycling with differentiation during neurogenesis. Development 138:4267–4277.

11. Florio M, Leto K, Muzio L, Tinterri A, Badaloni A, Croci L, Zordan P, Barili V, Albieri I, Guillemot F, Rossi F, Consalez GG. 2012. Neurogenin 2 regulates progenitor cell-cycle progression and Purkinje cell dendritogenesis in cerebellar development. Development 139:2308–2320.

12. Hindley C, Ali F, McDowell G, Cheng K, Jones A, Guillemot F, Philpott A. 2012. Post-translational modification of Ngn2 differentially affects transcription of distinct targets to regulate the balance between progenitor maintenance and differentiation. Development 139:1718–1723.

13. Bechard ME, Wright CVE. 2017. New ideas connecting the cell cycle and pancreatic endocrine-lineage specification. Cell Cycle 16:301–303.

14. Krentz NAJ, van Hoof D, Li Z, Watanabe A, Tang M, Nian C, German MS, Lynn FC. 2017. Phosphorylation of NEUROG3 Links Endocrine Differentiation to the Cell Cycle in Pancreatic Progenitors. Dev Cell 41:129–142.

15. Azzarelli R, Hurley C, Sznurkowska MK, Rulands S, Hardwick L, Gamper I, Ali F, McCracken L, Hindley C, McDuff F, Nestorowa S, Kemp R, Jones K, Göttgens B, Huch M, Evan G, Simons BD, Winton D, Philpott A. 2017. Multi-site Neurogenin3 Phosphorylation Controls Pancreatic Endocrine Differentiation. Dev Cell 274–286.

16. Sakaue-Sawano A, Kurokawa H, Morimura T, Hanyu A, Hama H, Osawa H, Kashiwagi S, Fukami K, Miyata T, Miyoshi H, Imamura T, Ogawa M, Masai H, Miyawaki A. 2008. Visualizing Spatiotemporal Dynamics of Multicellular Cell-Cycle Progression. Cell 132:487–498.

17. Chandler KJ, Chandler RL, Broeckelmann EM, Hou Y, Southard-Smith EM, Mortlock DP. 2007. Relevance of BAC transgene copy number in mice: Transgene copy number variation across multiple transgenic lines and correlations with transgene integrity and expression. Mamm Genome 18:693–708.

18. Abe T, Sakaue-Sawano A, Kiyonari H, Shioi G, Inoue K, Horiuchi T, Nakao K, Miyawaki A, Aizawa S, Fujimori T. 2013. Visualization of cell cycle in mouse embryos with Fucci2 reporter directed by Rosa26 promoter. Development 140:237–246.

19. Mellitzer G, Martín M, Sidhoum-Jenny M, Orvain C, Barths J, Seymour P a, Sander M, Gradwohl G. 2004. Pancreatic islet progenitor cells in Neurogenin 3-yellow fluorescent protein knock-add-on mice. Mol Endocrinol 18:2765–76.

20. Huang HP, Liu M, El-Hodiri HM, Chu K, Jamrich M, Tsai MJ. 2000. Regulation of the pancreatic islet-specific gene BETA2 (NeuroD) by Neurogenin 3. Mol Cell Biol 20:3292–3307.

21. Gasa R, Mrejen C, Lynn FC, Skewes-Cox P, Sanchez L, Yang KY, Lin CH, Gomis R, German MS. 2008. Induction of pancreatic islet cell differentiation by the Neurogenin-NeuroD cascade. Differentiation 76:381–391.

22. Horton RM, Cai Z, Ho SN, Pease LR. 2013. Gene splicing by overlap extension: Tailor-made genes using the polymerase chain reaction. Biotechniques 54:528–535.

23. Chintinne M, Stangé G, Denys B, In ‘T Veld P, Hellemans K, Pipeleers-Marichal M, Ling Z, Pipeleers D. 2010. Contribution of postnatally formed small beta cell aggregates to functional beta cell mass in adult rat pancreas. Diabetologia 53:2380–2388.

